# Selecting optimal support grids for super-resolution cryogenic correlated light and electron microscopy

**DOI:** 10.1101/2022.12.05.519127

**Authors:** Mart G. F. Last, Maarten W. Tuijtel, Lenard M. Voortman, Thomas H. Sharp

**Affiliations:** Department of Cell and Chemical Biology, Leiden University Medical Centre, 2300 RC Leiden, The Netherlands; Department of Molecular Sociology, Max Planck Institute of Biophysics, Max-von-Laue-Straße 3, 60438 Frankfurt am Main, Germany

## Abstract

Cryogenic transmission electron microscopy (cryo-TEM) and super-resolution fluorescence microscopy (FM) are two popular and ever improving methods for high-resolution imaging of biological samples. In recent years, the combination of these two techniques into one correlated workflow has gained attention as a promising route towards contextualizing and enriching cryo-TEM imagery. A problem that is often encountered in the combination of these methods is that of light-induced damage to the sample during fluorescence imaging that renders the sample structure unsuitable for TEM imaging. In this paper, we describe how absorption of light by TEM sample support grids leads to sample damage, and we systematically explore the importance of parameters of grid design. We explain how, by changing the grid geometry and materials, one can increase the maximum illumination power density in fluorescence microscopy by up to an order of magnitude, and demonstrate the significant improvements in super-resolution image quality that are enabled by the selection of support grids that are optimally suited for correlated microscopy.

## Introduction

Cryogenic transmission electron microscopy (cryo-TEM) can deliver structural information of unlabelled samples, such as proteins and nucleic acids^1,2^. Indeed, cryo-TEM can determine high-resolution protein structures even from crowded molecular environments, such as within cells. However, such *in situ* structural biology is currently only possible for common, readily-identifiable and large biomolecular complexes, such as ribosomes or proteasomes^3–5^. This is because it is challenging to identify specific structures of interest solely on the basis of charge density maps, especially in the crowded environment of the cell. The opposite is true in fluorescence microscopy (FM), where the spatial resolution is comparatively limited but specific labelling techniques enable the visualisation and identification of individual species of interest. These differences in resolution and labelling specificity make cryo-TEM and cryo-FM highly complementary techniques. The combination of the two techniques, commonly called ‘correlated cryogenic light and electron microscopy’ (cryo-CLEM), offers both the high-resolution spatial information acquired by cryo-TEM, as well as the specificity of labelling of FM. Despite recent advances in cryo-CLEM for structural biology^6–10^, combining and correlating these imaging modalities also introduces technical issues that must be addressed before the technique can be adopted more broadly.

A critical requirement in cryo-CLEM is the need to maintain sample integrity during the fluorescence imaging step, such that specimen quality is not compromised prior to cryo-TEM imaging. Exposure of the sample to high powered excitation light sources has been found to cause devitrification, melting and sublimation of the cryogenic sample^10,11^. The precise limit of this photon dose has yet to be determined, and reported values vary widely. For instance, Kaufmann et al. (2014)^12^ used an incident power of up to 2 kW/cm^2^ but did not report on devitrification, while Moser et al. (2019)^10^ claimed that an illumination density of around 200 W/cm^2^ was enough to cause obvious devitrification, while they found no evidence of devitrification at 100 W/cm^2^. Using a similar but less devitrification-prone holey carbon support film, Dahlberg et al. (2020)^8^ used no more than 50-100 W/cm^2^ to ensure that the sample remained vitrified, whilst Tuijtel et al. (2019)^13^ found a threshold for damage at ∼30 W/cm^2^ for similar grid types. Despite the lack of a consensus, it is clear that the intensity of excitation light that can be used is limited, that the choice of TEM grid support material is an important parameter^11^, and that sample thickness and use of cryoprotectants also affect the power limit^14^.

Here, we systematically explore the influence of sample support to determine how fluorescence imaging causes damage to vitrified samples. We demonstrate how the selection of suitable cryo-TEM supports and grids can alleviate this issue by comparing different support film material, film geometry, TEM grid materials, and grid mesh size. Using correlated imaging data collected on various types of grids, we conclude that a simple one-dimensional heat transport model with heat generated by absorption of light is sufficient to explain previous observations. This model also provides a guide for the selection of suitable support grids.

## Results

### Light-induced damage to cryogenic samples can be observed by light microscopy

Devitrification occurs when vitrified cryogenic samples are heated to more than approximately 136 K^15,16^. Above this temperature, amorphous ice crystallizes to form cubic ice^17^, or sublimates at even higher temperatures. The growth of such crystalline structures poses two problems for cryo-TEM imaging: i) the forces exerted during this phase transition can damage or destroy nearby structures of interest, and ii) the crystalline ice strongly scatters incident electrons, which obscures the TEM image.

The vitrification state of a sample is typically only observed once the sample is imaged in the TEM. Consequently, if a sample is devitrified during an earlier stage of the experiment, this will not be observed before performing the, often time-consuming, sample preparation process for cryo-TEM. In order to avoid this problem in cryo-CLEM experiments, and to be able to assess whether a sample had been exposed to excitation light of a power beyond the devitrification limit, we first tested whether it was possible to image (changes in) the vitrification state during the light microscopy step. Using a narrow band LED light source, we could acquire images of the light reflected by the ice and carbon in a sample. Since the layer of vitreous water in a sample for cryo-TEM is very thin (around 100 nm), the effects of thin-film interference are very pronounced in these reflected light images (**Fig. 1a**). By acquiring such reflected light images both before and after exposure of the sample to intense laser (488 nm) irradiation, we could directly image the effect of the exposure on the integrity of the sample (**Fig. 1b**). To confirm that these effects are indeed due to changes in the structure of the sample, we imaged the sample by cryo-TEM after laser illumination (**Fig. 1a,b**). The resulting cryo-CLEM image pairs show similar features. When a grid square is affected by light, the damage is typically most severe at the centre of the square, and less severe or even absent near the edges. Different regions on a single grid square thus show various degrees of damage. Cryo-CLEM imaging of individual holes in the carbon support film of over-exposed grid squares revealed several discernible effects. In order of decreasing severity, regions can be entirely destroyed, devitrified to a varying extent, or unaffected (**Fig. 1b**). Each of these stages presents differently and can be recognized in reflected light microscopy images: the absence of ice results in reduced reflectivity (panel *i*), the presence of large particles of devitrified ice in the middle of a hole is marked by a bright reflective centre (panels *ii & iii*), thickening or crystallization of ice around the edges of a hole results in increased reflection around the edges and a concomitant decrease in reflection of the adjacent carbon (panels *ii-v*), and the deposition of small particles increase reflectivity of the hole (panels *iv & v*). The rightmost panel (panel *vi*) shows a hole containing vitreous water. These data indicate that the structural changes due to laser illumination can be observed during the initial light microscopy step of a cryo-CLEM experiment.

**Figure 1.**
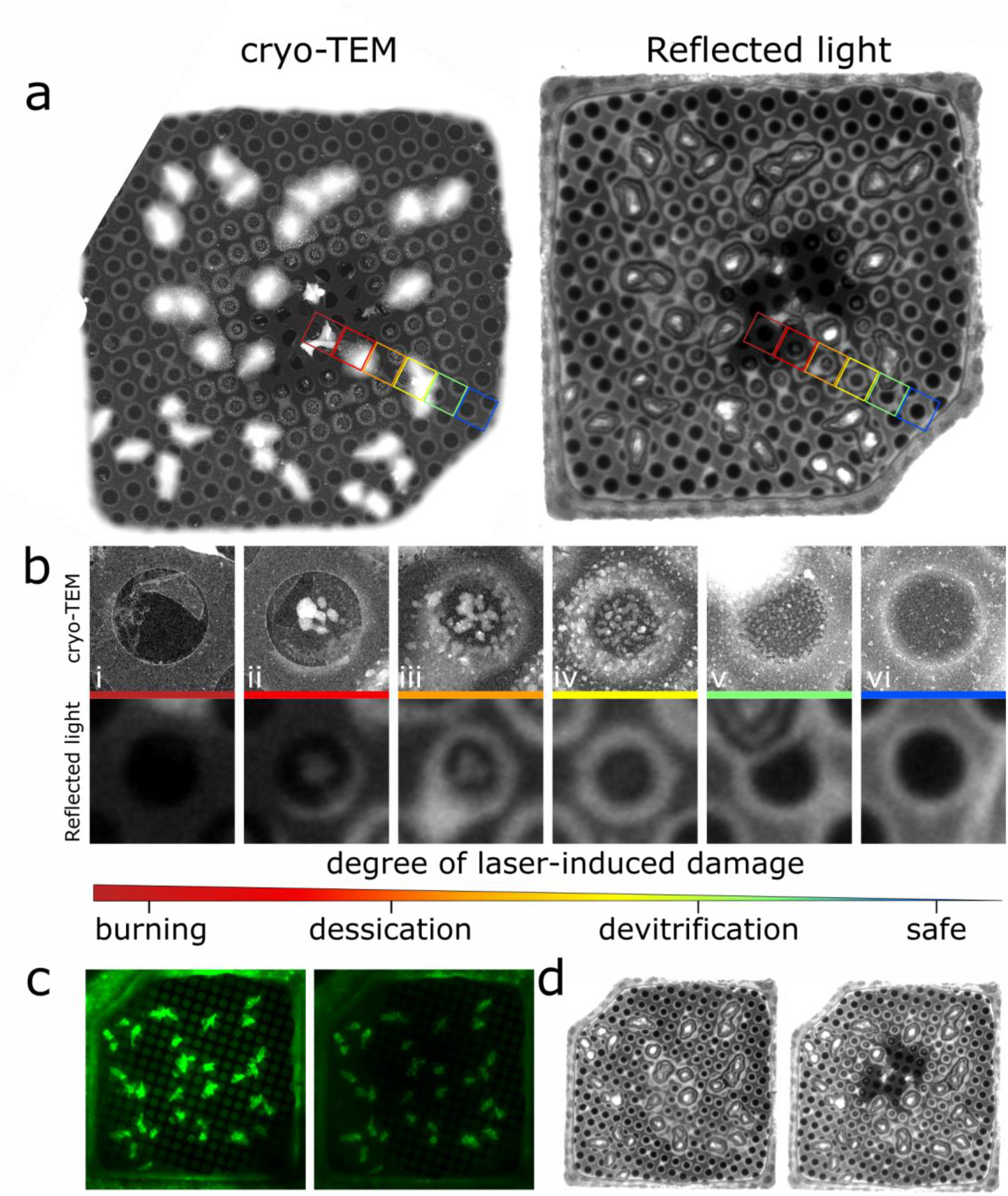
Reflected light and fluorescence microscopy images of cryo-samples reveal the effect of high-power illumination. **a)** Cryo-TEM and reflected light images of the same region of a holey carbon TEM grid after exposure to high-power laser illumination. Coloured squares indicate the position of holes shown in the below image panel. Note that the contrast in the cryo-TEM images was inverted in order to facilitate image comparison. **b)** Single holes in the carbon film imaged at higher magnification cryo-TEM (top row) or cryo-LM (reflected light, bottom row), revealing various degrees of laser-induced damage (i-vi). **c)** Typical fluorescence images of a grid square before (left) and after (right) exposure to intense laser light. The images are presented at the same contrast/brightness settings. **d)** Typical reflected light images of a grid square (same region as in panels **a** and **c**) before (left) and after (right) illumination.

Imaging cryo-TEM grids using reflected light therefore provides a simple but useful method of screening sample quality prior to cryo-TEM imaging, which can also be used to determine the maximum photon illumination power density of a sample. By imaging samples containing a low concentration of EGFP (10 μM) using both reflected light and fluorescence, we could screen many different types of TEM grids to find their maximum illumination power density. A comparison of images of a region before and after illumination (**Fig. 1c,d**) is sufficient to establish whether a chosen laser power was above or below the devitrification limit for that particular sample support.

### Temperature changes in grid squares is governed by a 1D steady state heat equation

When grids were damaged by light, this damage was typically most apparent at the centre of the grid, while regions close to the grid bars were often not affected (**Fig. 1a,b**). We also observed that the damage occurred near-instantaneously (within tens of milliseconds) when the incident laser power was raised above the devitrification limit (**Fig. 2a**). Exposure of a grid square to 56 W/cm^2^ for 25 seconds revealed no discernible damage (panel *i*), yet imaging at 254 W/cm^2^ for just 5 seconds resulted in clear devitrification (panel *ii*). This damage occurred withing 10 ms of illumination above the devitrification limit (panels *iii & iv*). When, as in this case, the support film is torn or otherwise damaged, the effects of exposure to intense light can be seen near instantaneously. This indicates that it is not the total dose that governs the degree of damage, but rather the dose rate; i.e., the incident power density, as previously observed ^13^.

**Figure 2.**
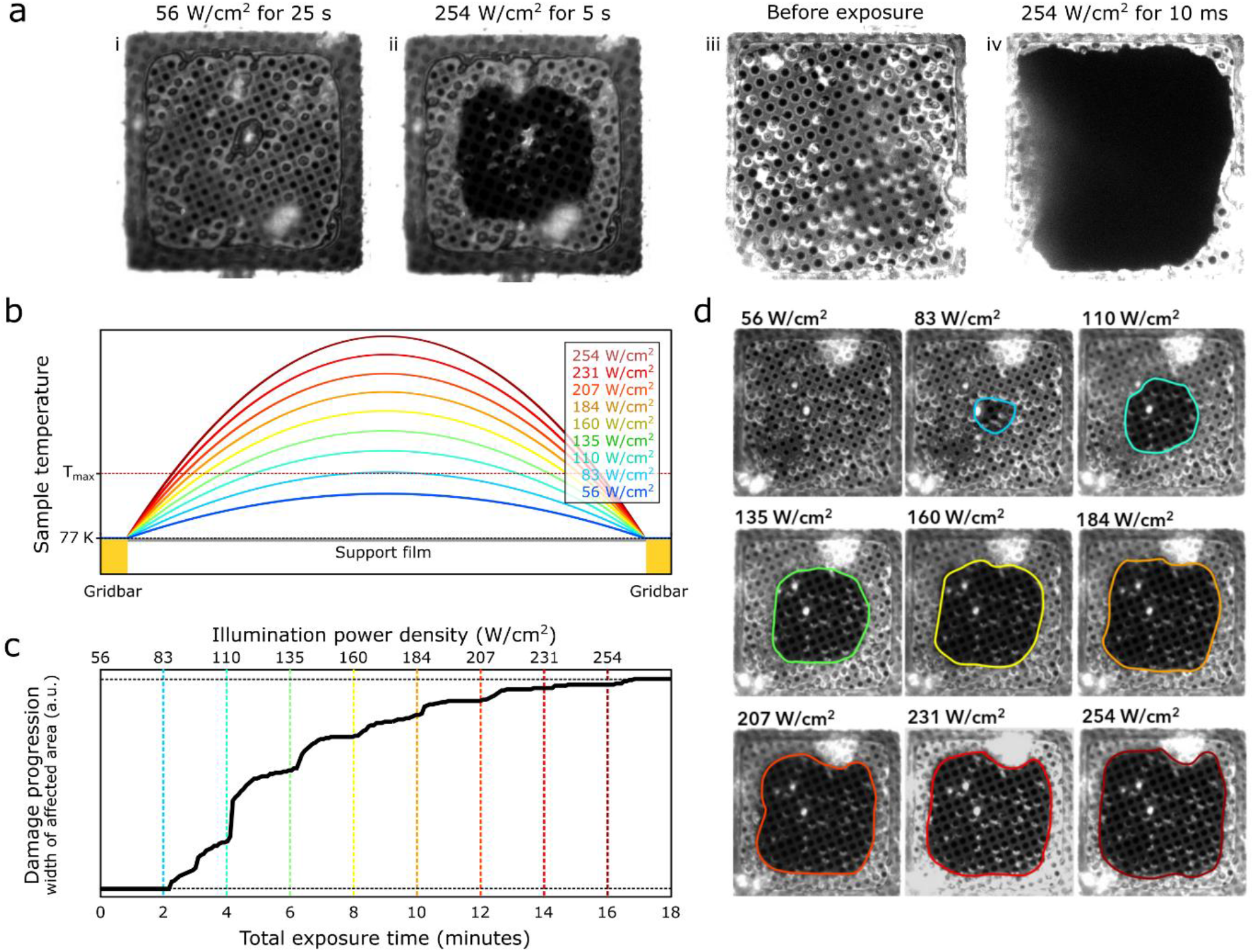
A steady-state heat transport model explains how heat generated by absorption of light damages the sample. **a)** The same grid square is shown after: i) 25 seconds exposure to 56 W/cm^2^, ii) 5 seconds exposure to 254 W/cm^2^. A separate grid square is shown before (iii) and after (iv) 10 ms exposure to 254 W/cm^2^. **b)** Steady-state temperature profiles, simulated according to the model described in the main text. At higher incident powers, a larger fraction of the area of a grid square reaches a temperature above T_max_, a temperature where damage begins to occur. **c)** A plot of the damage progression over time, with the illumination power density setting indicated above. **d)** Reflected light images of a grid square after exposure to increasing illumination power densities. The size of the affected spot is seen to grow, in accordance with the model depicted in panel b.

A simple one-dimensional, steady-state heat transport model can qualitatively explain these observations. We posit that the absorption of light by the support film causes heating of the sample. This film is in thermal contact with the metal grid bars, which are held at a constant temperature of 77 K by the liquid nitrogen-cooled sample stage. Besides this conductive cooling, the sample is also convectively cooled by the surrounding cold nitrogen gas. Since we expect convective cooling to be of lesser importance than conductive cooling, and that convection will affect different grid types equally and can therefore not explain observed differences, we do not include this term in our model.

The temperature profile along a line through the centre of a grid square can thus be found by solving the heat equation with constant boundary conditions and a source term dependent on both the illumination power density and the film absorptivity (**Fig. 2b**). We also make the assumption that the thermal mass is very low in comparison to the used illumination power, and thus consider the system to be in steady state. The resulting equation and boundary conditions are as follows:

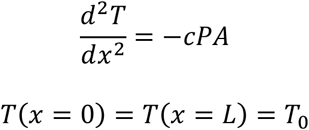

Here, *T*(*x*) is the temperature at a position 0 < *x* < *L* along a line through the centre of a gridsquare, *c* is the combined thermal conductivity of the support film and the vitreous sample, *L* is the size of a grid square (i.e. the distance between grid bars), *P* is the illumination power density, and *A* the support film absorptivity. The result is a qualitative relation between the sample temperature, the grid geometry and absorptivity, and the illumination power density (Fig. 2a):

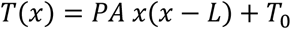

This result explains why light-induced damage first occurs in the centre of a grid square. Closer to a grid bar, the heat-sink effect of the cooled metal is efficient enough to keep the sample below the devitrification limit even at higher incident light powers, while the centre of a square (being most distant from the heat sink) reaches the highest temperature and as such will be affected at lower incident light levels than the outer edges of a grid square.

This model can be tested by monitoring the effect of exposure to laser light on a sample. By exposing the sample to an initially low powered laser, waiting for two minutes before stepping up the laser power to a higher value, and identifying the size of the damaged region on a grid square after each step, a damage front can be observed that progresses outward from the centre of the grid square toward the grid bars (**Fig. 2c,d**). The reason for this stepwise progression is as follows: when the laser power is increased, the temperature of the grid increases accordingly, causing a larger region of the grid to reach a temperature above the damage threshold. In these experiments it was also apparent that the heating of the sample can occur on a short timescale: the damaged region rapidly grows within the first few seconds of exposure, and then halts until the laser power is increased further.

A consequence of the steady-state heating behaviour of the grids is that the light-induced damage is dose independent. Larger total doses of light can be delivered to the sample with no effect on the vitrification state, as long as the dose rate does not exceed a certain critical value. Conversely, even a small dose of light can be detrimental when it is administered in an instant. For example, just 10 ms of exposure to 254 W/cm^2^ of light can be sufficient to completely destroy a grid square (**Fig. 2a**).

### The maximum illumination power density is not absolute

A well-defined threshold for the illumination power density would be useful for the design of cryogenic fluorescence imaging experiments. In practice it is difficult to define such a specific threshold, because the intensity of illumination that a grid can sustain depends not just on the support film material and grid geometry, but also on the thickness of the ice, integrity of the support film, the presence of cryoprotectants such as glycerol or cellular material, and the composition of the sample itself^14^.

Furthermore, different cryostages may present different thermal conditions. Nevertheless, although the resulting maximum value may differ, the general conclusions about the importance of grid geometry and material will be the same.

To estimate the illumination threshold, we therefore used exemplary samples consisting of a low concentration of fluorescent protein (10 μM EGFP) in phosphate buffered saline (PBS) deposited onto grids and vitrified by plunge-freezing. Having a thin layer of vitreous ice and lacking cryoprotectants, we could use these samples to isolate the effects of different support films and grid geometries. We then used a system where we classified grid squares after illumination as being in one of four damaged states of increasing severity, on the basis of both reflected light and fluorescence images (**Fig. 3**). These are 1) no effect or uniform bleaching alone, 2) increased bleaching in the centre of the grid square and concomitant, slight, but observable effect in the reflection images, 3) notable effect on both fluorescence and reflection images in the centre of the grid square, and 4) notable effect on both fluorescence and reflection images across the entire grid square. Clearly, states 3 and 4 are incompatible with subsequent cryo-TEM imaging, while state 1 is acceptable. State 2 appeared to represent the onset of light-induced changes to the sample, with a slight but noticeable effect on the ice structure and fluorescence uniformity that provided a consistent and reliable model to assess the effect of illumination upon cryosamples. Therefore, we selected state 2 as the indicator for the onset of light-induced changes to the sample, and consequently the power density that caused these effects as the maximum illumination power density for that grid type.

**Fig. 3.**
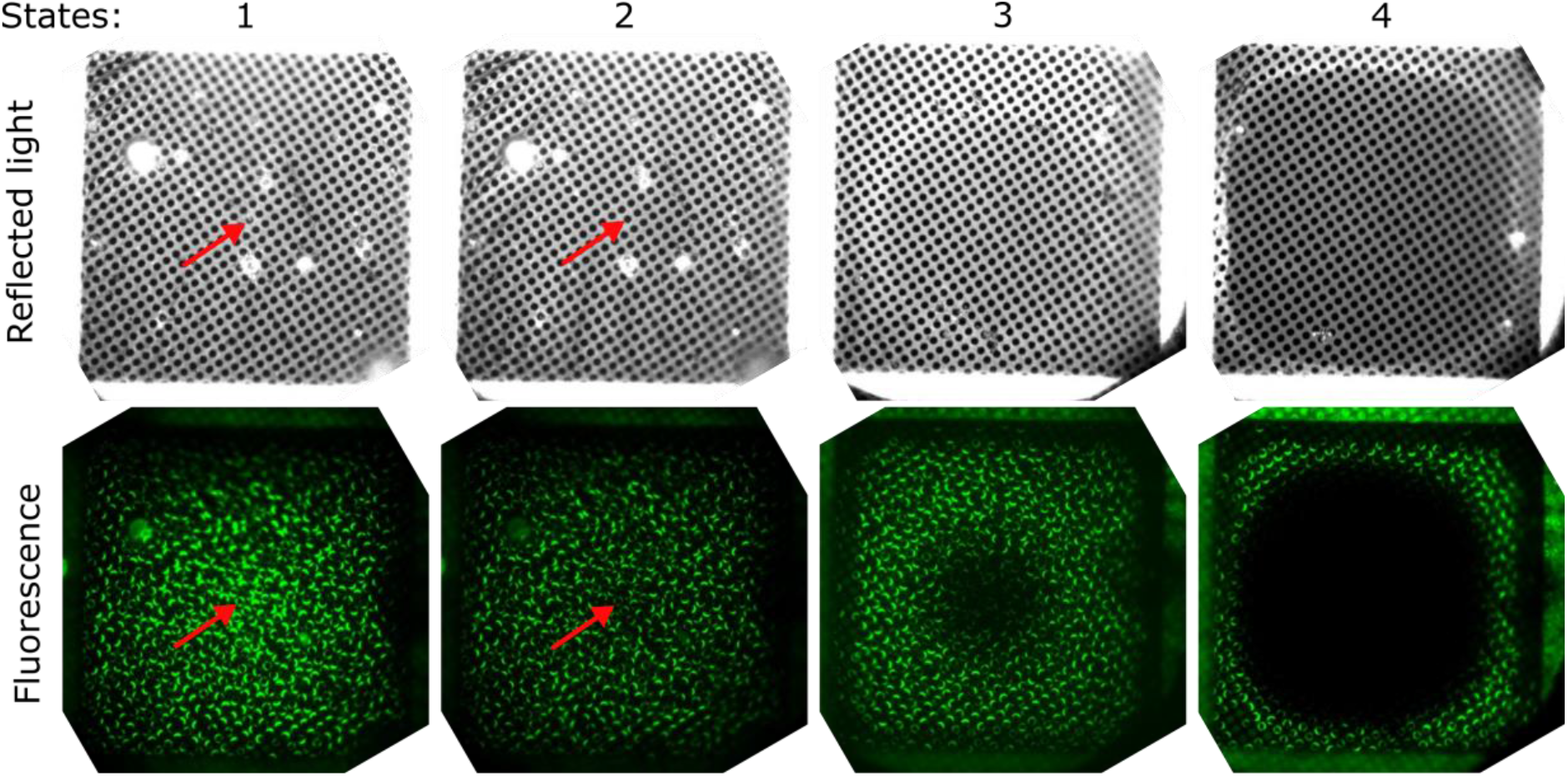
Reflected and fluorescence light images of post-illumination states showing degrees of devitrification. **State 1**: No visible differences between pre- and post-illumination images of the grid square. **State 2**: the same grid square as shown in 1 illuminated at a higher intensity shows barely discernable bleaching of fluorescent proteins in the centre of the grid square and concomitant slight changes observed in the reflected light images, such as disappearance of small particles in the centre of the grid square, and a reconfiguration of the stress pattern in the support film – compare the images of state 1 and 2 at positions indicated by red arrowheads. **State 3:** notable effects in both the reflected light and fluorescence images. Significantly increased bleaching of fluorescent protein near the centre of the grid square, and a marked darkening seen in the reflected light images. Effects are pronounced, but not complete. **State 4:** complete desiccation of the grid over a large fraction of the area is visible in reflected light images as well as near-total bleaching of fluorescence.

### Selecting grids that enable higher illumination powers

The relation between the temperature profile across a grid square and the illumination power density, film absorptivity and grid geometry suggests multiple ways of reducing the temperature increase upon illumination and increasing the devitrification limit: using less absorptive materials, different film geometries, thinner films, or grids with a geometry that enhances heat transport to the grid bars. By determining the maximum illumination power densities for various grids by using the criteria described above, we were able to isolate the effect of each of these four strategies. A table of the results is presented and discussed below (**Table 1**).

**Table 1.**
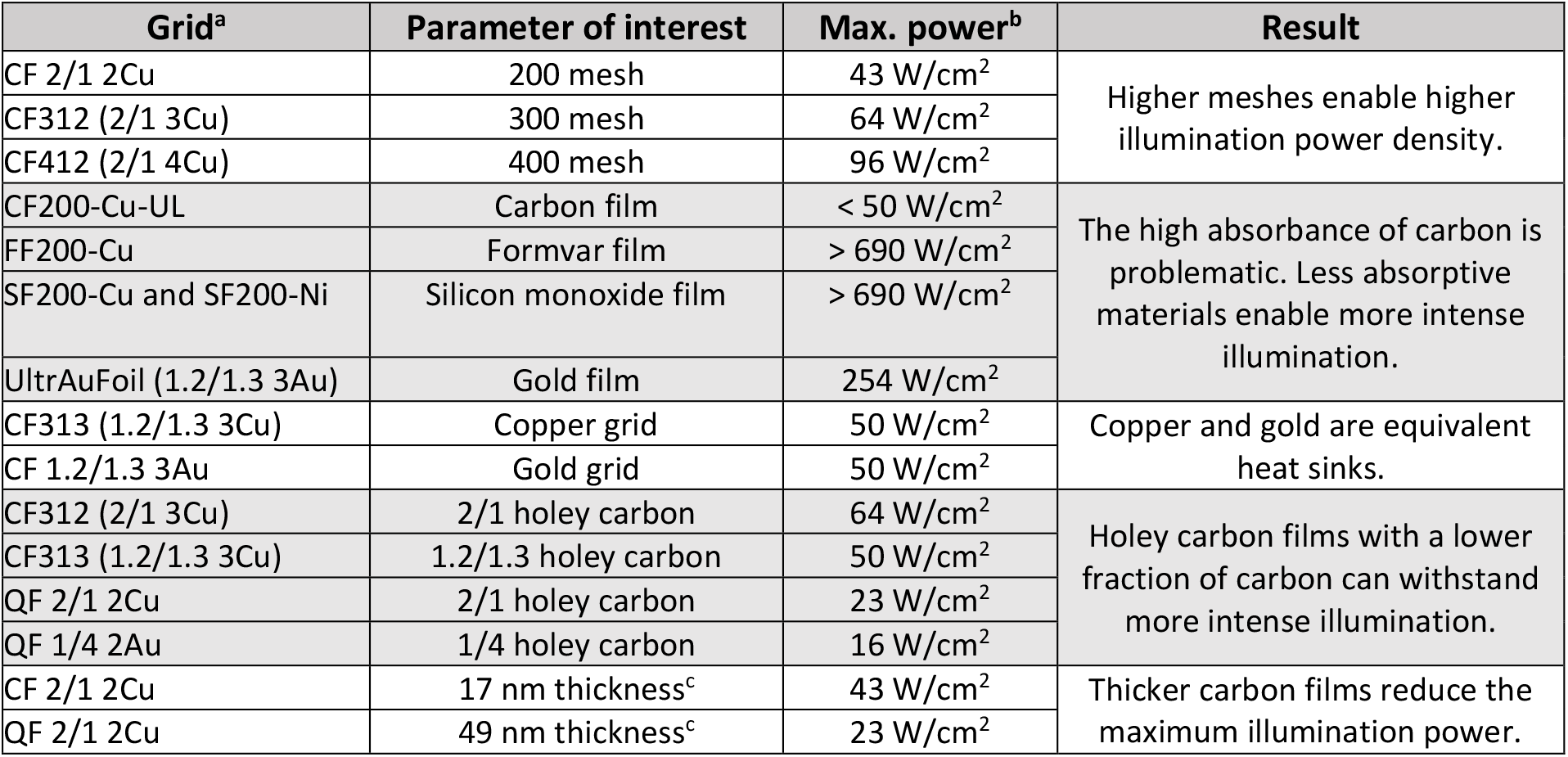
overview of maximum illumination power densities (at 488 nm) for various types of TEM grids. ^a^All grids other than the Quantifoil grids were ordered from EMSDiasum and are named using the supplier product name with an optional clarification in brackets. ‘CF’ indicates CFlat, ‘QF’ Quantifoil. For holey carbon grids, the hole size and spacing are indicated as, e.g., 2/1 (for 2 μm holes with 1 μm spacing.) The grid metal is indicated as Cu/Au (copper, gold), alongside a number that signifies the mesh (2, 3 or 4, for 200, 300, or 400 mesh. Corresponding grid square sizes are 90, 58, and 37 μm, respectively.) ^b^Due to the low absorptivity of both formvar and silicon monoxide, we were not able to reach the maximum illumination power density of these grids. The highest power density reached for these experiments was 690 W/cm^2^ (combining 367 W/cm^2^ of 488 nm and 323 W/cm^2^ of 405 nm). ^c^See reference^18^.

These results imply a number of interesting points of consideration in the selection of optimal grids for cryogenic fluorescence microscopy regarding film and grid material, film and grid geometry, and film thickness, which we discuss below.

### Film material

The most important factor that determines the maximum illumination power density is the material of the support film. While carbon support films are commonly used for cryogenic fluorescence microscopy, we found that replacing the carbon with formvar or silicon monoxide, or to a lesser extent gold film, drastically increases the power limit. This has previously been realized by Liu et al. (2015), who also noted that formvar autofluorescence posed a problem for super-resolution cryogenic microscopy^11^. Likewise, we observed that commercial formvar (FF, formvar film) and silicon monoxide (SF, silicon film) grids exhibited very bright, blinking autofluorescence within the spectral range of common green fluorescent proteins, which renders these grids unsuitable for single molecule localization fluorescence microscopy, but may find utility for other imaging methods. Various attempts at cleaning the SF grids with chloroform, acetone, or by brief etching with a mixture of hydrogen peroxide and ammonium hydroxide, all failed to remove the fluorescent material. Thus, although FF and SF appear to be promising materials for TEM grids for cryo-CLEM, the lack of commercially available clean grids (see Methods) means the use of these films is currently impractical.

Grids with a pure, 50 nm thick, gold film (UltrAuFoil 1.2/1.3 Au 300, Quantifoil) also sustain higher illumination powers, although the limit is lower than with the FF and SF grids. We attribute this to the relatively high absorption of gold at 488 nm compared to silicon monoxide^19,20^. Besides enabling an illumination power more than five times the limit of a comparable carbon-film grid, gold film also has advantages in TEM such as reduced beam-induced sample motion^21^ and superior flatness, is commercially available, and has no background fluorescence, and as such is our film material of choice for high-power fluorescence imaging of cryosamples.

### Grid material

In another test, we compared grids with the same support film (1.2/1.3 holey carbon, see next section) and mesh (300) but that differed in the type of metal base, using either copper or gold. Copper grids are more affordable than gold grids, but are not suited for cell culture due to copper toxicity. Since we used copper grids for most of our experiments, we wanted to determine whether the type of metal had an effect on the maximum power density. As expected, we were unable to measure a difference in the maximum illumination power density between the types of metal, indicating that both copper and gold are sufficiently thermally conductive for dissipation of heat through the metal not to be the limiting factor.

### Film geometry

Holey carbon is a type of discontinuous support material, comprising regularly interspaced holes of specified size in a thin carbon film. These grids are commonly used for cryo-TEM, and are widely available with various configurations of hole size and spacing, such as 2/1 (2 μm holes and 1 μm spacing), 1/4, and 1.2/1.3. In principle, each of these is suited for growing cells and cryo-CLEM, but an important difference is that the different geometries also differ in the fraction of the film that is composed of carbon. A film with a large fraction of the area consisting of carbon, such as is the case for 1/4 where 97% of the area is carbon, absorbs more light per unit area than a low carbon fraction grid, such as 2/1, where 65% of the area is carbon.

We were interested in whether films with lower carbon fractions would enable higher illumination power, owing to the reduced absorption per area. To test this, we compared two pairs of grids: CFlat 2/1 and 1.2/1.3 (82% carbon) on 300 mesh copper, and Quantifoil 2/1 and 1/4 on 200 mesh gold (for 2/1) or copper (1/4). We expected the maximum power to be inversely proportional to the carbon area fraction, or, in other words, that the product of the carbon fraction *η*_*c*_ and the maximum power density *P*_*max*_ would be constant with other parameters (e.g. mesh) unchanged: *η*_*c*_*P*_*max*_ = *C*. The result showed that this was indeed the case: for the CFlat set, the value for *C* differs by ∼2%, while the difference is ∼3% for the pair of Quantifoil grids. Holey carbon films with larger holes and smaller spacings are therefore better suited for cryogenic fluorescence microscopy.

### Grid geometry

The temperature profile across a grid square is dependent on the size of the grid square, with the temperature in the centre reaching significantly higher values than near the grid bars. Grids with different meshes (number of grid squares per inch), such as 200, 300, or 400 mesh (corresponding to grid square sizes of 90, 58, and 37 μm, respectively) should be able to withstand different illumination power densities. By comparing 2/1 holey carbon grids on each of these three different mesh grids, we confirmed that large grid squares (e.g. 200 mesh, 43 W/cm^2^ maximum) are more susceptible to light-induced devitrification than small grid squares (e.g. 400 mesh, 96 W/cm^2^ maximum). To maximize the viable light intensity, higher mesh numbers can be used.

A downside of the use of high mesh grids is that in cryo-electron tomography, regions of the sample close to the metal grid bars are blocked from view by the grid bars when imaging at a high tilt angle. One solution to this problem could be to use grids with a rectangular mesh, where a high mesh number in one direction ensures efficient cooling of the sample, while a low mesh number in the other direction enables TEM imaging at high tilt angle, provided that the grid is properly oriented relative to the tilt axis of the TEM.

### Film thickness

Because the amorphous carbon used for TEM grids has a relatively low thermal conductivity (∼0.2-0.4 Wm^-1^K^-1^)^22^ compared to vitreous water (∼1.5 Wm^-1^K^-1^)^23,24^, we expected that grids with a thinner film of carbon could be safely imaged at higher incident power than grids coated with a thick carbon layer.

CFlat and Quantifoil are two commonly used holey carbon films that differ significantly in thickness, CFlat (∼17 nm) being approximately 2.5 times thinner than Quantifoil (∼45 nm)^25^. When comparing two grids that differ in no respect other than the make (200 mesh copper 2/1 holey carbon) we observed that the sample on the thicker Quantifoil film was damaged at a significantly lower (∼53%) incident power density than the CFlat film. Thus, when using carbon grids, high illumination power densities are enabled by thinner support films.

### Super-resolution fluorescence cryomicroscopy with a high illumination power density and without devitrification

To show the difference that the selection of a support grid optimally suited for high-powered illumination fluorescence imaging and subsequent cryo-TEM can make, we cultured human osteosarcoma (U2OS) cells expressing rsEGFP2-MAP2, a fusion of the green fluorescent protein rsEGFP2 that is known to efficiently photoswitch between bright and dark states at cryogenic temperature^13^, and MAP2, a microtubule binding protein, on 1.2/1.3 holey-gold 300 mesh gold grids (1.2/1.3 UltrAuFoil, Quantifoil). These grids were then imaged by fluorescence microscopy using 186.5 W/cm^2^ of 488 nm light – a value far exceeding the limit previously found for any type of carbon grid, but below the limit for gold film (we decided to operate laser at ∼70% of the previously determined maximum). To illustrate the advantage of using higher laser powers, we imaged other regions of these same samples using a lower laser power (34.2 W/cm^2^), which was the maximum safe laser power that we determined for CFlat 1.2/1.3 holey carbon 300 mesh gold grids, the most similar type of holey carbon grid. Since rsEGFP2 is a negatively photoswitchable fluorescent protein and it is common in single-molecule localization microscopy experiments to deplete fluorescence prior to acquiring data for super-resolution fluorescence microscopy, a comparison of the photo-deactivation kinetics under these two illumination conditions is a useful measure to determine how a higher illumination power affects data acquisition. We found that with the high-power illumination settings, the half-time of fluorescence decay was approximately eight times shorter (31 versus 248 seconds) than with the low power settings (**Fig. 4a**).

**Figure 4.**
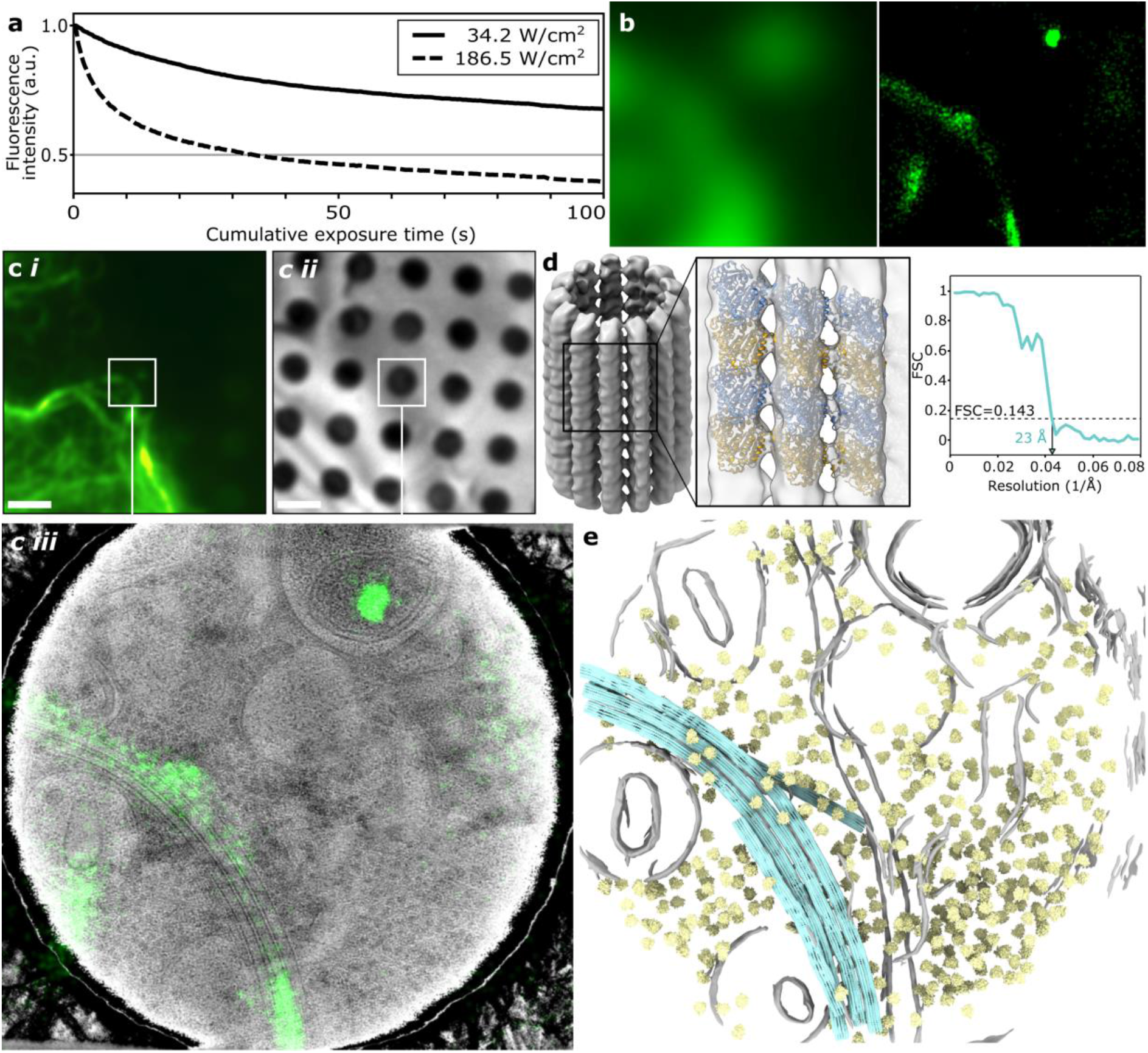
Correlated super-resolution fluorescence cryo-microscopy and cryo-TEM using gold film TEM grids. **a)** Photo-deactivation and -bleaching of rsEGFP2 under exposure to either 34.2 W/cm^2^ (solid line) or 186.5 W/cm^2^ (dashed line) on 1.2/1.3 CFlat holey carbon 300 mesh and 1.2/1.3 UltrAuFoil holey gold 300 mesh grids, respectively. **b)** Resolution improvement from widefield (left) to super-resolved reconstruction (right). **c)** Super-resolved reconstruction (**i**) and reflected light image (**ii**) correlated (**iii**) showing microtubule cytoskeleton. **d)** Sub-tomogram average (left) of microtubules extracted from cellular tomograms at regions located using super-resolution imaging with fitted model (middle). The map reached 23 Å resolution with no symmetry applied (blue). **e)** Fitting the microtubule map back into the tomogram, together with identified ribosome locations (yellow) and membranes (grey). Scale bars are 200 nm (b) and 2 μm (c, d).

Higher laser powers thus enable super-resolution data acquisition in a shorter time. Here, using a cumulative exposure time of only 100 seconds (1000 frames with 100 ms exposure time per frame), we were able to generate a super-resolution reconstruction containing hundreds of localizations within an area of ∼1 μm^2^ (**Fig. 4b,c**). After light imaging, we performed cryo-electron tomography on the same regions of interest in order to determine whether the structure of the sample was indeed still intact (**Fig. 4c**); no devitrification or other artefacts were seen in cryo-electron tomograms acquired in regions of the sample that were exposed to laser light, and the visibility of structures such as lipid bilayers, microtubules and ribosomes also indicates that the sample structure was not affected by illumination. In keeping with the earlier maximum illumination power density measurements with EGFP, bleaching of the fluorescent signal was uniform, and no changes could be observed in the reflected light images.

Finally, we performed sub-tomogram averaging on the microtubules found in 7 of tomograms for which we had also acquired super-resolution data. This resulted in a structure at 23 Å resolution that could be fitted back into the tomographic volumes (**Fig. 4d,e**).

## Discussion

Identifying the ideal sample supports for cryogenic samples on TEM grids is crucial for effective data collection. Here, we systematically explored various factors that determine to what extent cryogenic samples on TEM grids are affected by exposure to high-intensity excitation light. We find that the most important factor that determines the illumination power density of a sample is the material that the support film of the grid consists of (**Table 1**). Furthermore, other parameters, such as the grid geometry, support film geometry, and the film thickness, can additionally help to increase the maximum illumination power density to raise the devitrification limit.

The problem of light-induced devitrification of samples for cryo-CLEM can be solved in different ways. Examples are the use of cryoprotectants^14^ or by performing experiments at exceedingly low temperatures, e.g. by using liquid helium as the coolant.^7^ Such approaches require either the addition of possibly undesirable chemicals to the sample, or infrastructure for working with liquid helium and the use of custom cryogenic sample stages. Because grid selection poses fewer restrictions on the sample itself or the cryogenic stage that is used, we believe that our findings may be broadly useful for the design and use of cryogenic fluorescence microscopy. Based on our results, we identify two avenues towards increasing the maximum illumination power density for cryogenic (super-resolution) fluorescence microscopy.

The first approach may be valuable when the use of carbon for the support material is required, in which case a high mesh grid (i.e., small grid squares) combined with a thin carbon film with a low carbon fraction (i.e., large holes with small separation) are ideal. Furthermore, the use of grids with a non-square geometry designed to minimize the distance between regions on the support film and the heat-sinking metal would be advantageous. Many suppliers offer rectangular meshes where the high mesh number is 400, but a more significant gain in the maximum power density would be achieved at even higher mesh values, when the spacing between grid bars is much less than the 400-mesh 37 μm.

The second option is to replace the carbon as the material for the support film. Recently, metal support films have proven useful to inhibit devitrification for cryo-light microscopy of bacterial samples^26^. Here we extend this to additional support films for intracellular mammalian proteins. Specifically, gold films or other novel support films, such as ultraclean plastic or silicon oxide, with low-absorption and low background fluorescence. In experiments where it is not necessary to maximize the excitation power density, the low absorptivity of gold film enables the use of grid geometries that would otherwise be inefficiently cooling, such as 200 or even lower mesh grids. These grids would provide able area for cells to grow at a distance from the metal grid, yielding more regions on the sample that are also suited for high-tilt imaging in cryo-TEM.

An important parameter that determines the maximum illumination power density of the sample that is typically less controlled than the previous parameters is the thickness of the sample. As noted before, the thermal conductivity of amorphous ice is relatively high, and as such the sample itself presents the best avenue for heat transport towards the cold metal of the grid bars. Cryo-TEM requires that the sample be sufficiently thin as to be electron-transparent (i.e. at most a few hundred nanometers). However, when performing fluorescence imaging of whole cells grown on grids, regions adjacent to a thin region of interest can be much thicker than that, and as such may locally enable a significantly higher illumination power density. This buffering effect could potentially be especially pronounced when the region of interest itself also has a high thickness, for example in cases when cells are imaged by fluorescence microscopy prior to thinning of the sample by focused ion beam (FIB) milling^27^. However, in our experiments we employed grids hosting a very thin layer of sample, and we therefore note that the measured values for the maximum illumination power density of the various grid types may underestimate the actual maximum power densities achievable when performing cryogenic fluorescence imaging on thicker samples, such as cellular material. Comparisons between the maximum illumination densities of different grid types hosting identical samples, however, will still be valid.

In conclusion, we found that by using gold-film grids, we could drastically improve the devitrification limit by avoiding heating of the sample, and thus increase the maximum illumination power density used in our super-resolution fluorescence imaging experiments by more than a factor of five in comparison to commonly used carbon grids. This enabled faster data acquisition (a few minutes per region of interest, compared to tens of minutes in earlier experiments^28^), and the localization of hundreds of fluorescent proteins within the area of a single cryo-electron tomogram.

## Methods

### Super-resolution fluorescence cryomicroscopy

All cryogenic fluorescence microscopy imaging was performed using a custom wide-field fluorescence microscope in combination with a commercial cryogenic sample stage (Linkam Scientific, CMS196v3). Fluorescence of EGFP was excited with a 488 nm laser (Omicron, LuxX 488-100) and collected with a CMOS camera (pco.edge 4.2), using a 0.9 NA air immersion objective lens (Nikon, CFI TU Plan Apo 100x/0.9NA), a 495 nm edge dichroic mirror (Semrock, FF495-Di03), a band-pass emission filter (Semrock, FF01-540/80), and a 200 mm tube lens (Applied Scientific Instruments, C60-TUBE-B). Light from an LED source (Omicron, LedHUB 530 nm) is coupled into the laser illumination path by use of a low-reflectivity beam splitter glass. The illumination power density of the microscope was determined by measuring the total laser power incident on the sample plane (Thorlabs, PM200) and dividing this value by the area of the illuminated field (119 μm in diameter, visible in the microscope field of view.) This measurement was performed for every laser power setting (between 1-100%) that was used in the experiments. Super-resolution reconstructions of time-lapse videos of rsEGFP2 fluorescence were prepared using ImageJ^29^ plugins ThunderSTORM^30^ and TurboReg^31^.

### Sample preparation

To prepare the samples used to determine the maximum illumination power densities, grids were plunge-frozen in liquid ethane (-183 °C) using a Leica EM GP operated at 12 °C and 50% humidity (EGFP samples). 3 μL of sample (10 μM EGFP in PBS) was deposited onto glow-discharged grids (Electron Microscopy Science and Quantifoil – see Table 1), prior to blotting the grids from the sample-side for 3 seconds.

The samples with U2OS cells were prepared as follows: grids (1.2/1.3 UltrAuFoil, Quantifoil) were glow-discharged, submerged in sterile PBS in a 1 ml cell culture dish, and sterilized by UV-irradiation in a cell culture hood for 30 minutes. After sterilization, the PBS was removed and the dish filled with 1 mL of Dulbecco’s modified eagle medium (DMEM) supplemented with penicillin/streptomycin and fetal bovine serum. U2OS cells stably expressing rsEGFP2-MAP2 (generated by transfection of U2OS cells with a plasmid encoding rsEGFP2-MAP2 and neomycin resistance, followed by selection with neomycin) were then seeded (100000 cells per 35 mm culture dish) and incubated at 37 °C and 5% CO_2_ for 16-24 hours, after which the grids were plunge-frozen.

### Cryogenic transmission electron microscopy and correlation

After fluorescence imaging, grids were either stored in liquid nitrogen or immediately transferred to a Gatan 626 cryogenic sample holder and inserted into a Tecnai T12 transmission electron microscope (FEI) operated at 120 kV and equipped with a OneView CMOS detector (Gatan). All images were recorded in low-dose mode to avoid damage to the sample that could otherwise have been interpreted as due to exposure to laser light. The data presented in Figure 3b is an exception: electron tomography on the cellular samples was done using a Talos Arctica 200 kV cryo-TEM (Thermo Fisher Scientific) equipped with a K3 direct electron detector and BioContinuum energy filter (Gatan). Cryo-TEM and cryo-fluorescence images were correlated manually by matching features visible in both images and adjusting the transform of the TEM images such that these features overlapped.

### Subtomogram analysis

Cryo-electron tilt series were aligned using fiducialless patch tracking in IMOD^32^. The aligned tilt series were imported into EMAN2^33^, which was then used to reconstruct tomographic volumes binned by 4 with no additional alignment. Tomograms were segmented using in-built neural network procedures to identify membranous organelles. To reconstruct microtubules, The path of microtubuli in 7 tomograms were manually traced within EMAN2, yielding a total of 24 filaments. Boxes of 180^3^ pixels were extracted along each filament with an overlap of 75% and an Ångstrom/pixel value of 2.75, yielding 1,599 particles. These were aligned within EMAN2 to an initial model of 13 protofilaments filtered to 50 Å resolution using a helical pseudosymmetry characteristic of microtubules; H13:1:27.6923:3.4, corresponding to a 13 protofilament tubule with an nstart of 1, a subunit rotation of 360/13 and an axial rise of ∼0.9 nm Å (3.4 pixels). The resulting average was place back into the tomograms using e2spt_mapptclstotomo.py within EMAN2. Ribosomes were picked from the tomograms using the neural network within EMAN2, which were aligned to an 8 Å lowpass-filtered map of the human 80S ribosome (EM database deposition code EMD-2938) over 8 iterations, and then placed back into the tomograms using e2spt_mapptclstotomo.py as above.

## Author Contributions

M.G.F.L. performed all data collection and light microscopy image analysis. M.W.T. and L.M.V. contributed to image analysis. M.G.F.L. and T.H.S. performed cryoEM image analysis. L.M.V. and T.H.S. supervised the project. All authors contributed to and reviewed the manuscript.

## Acknowledgments

We thank L. Janssen for his help in generating the rsEGFP2-MAP2 expressing U2OS cell line. This research was supported by the following grants to THS: European Research Council Grant 759517; The Netherlands Organization for Scientific Research Grants OCENW.KLEIN.291 and VI.Vidi.193.014.

